# SNCA Deletion Induced Aberrant Projection of Olfactory Sensory Neurons via NCK2-EphA4 Pathway

**DOI:** 10.1101/2022.10.25.513708

**Authors:** Jing Ren, Chao Wu, Jingjing Yue, Mengxia Zeng, Mingqin Qu, Ning Chen, Ge Gao, Yuwen Jiang, Jing Liu, Baoyang Hu, Hui Yang, Yin Jiang, Fangang Meng, Jianguo Zhang, Ling-ling Lu

**Affiliations:** Department of Neurobiology, Capital Medical University, Beijing 100069, China; Key Laboratory for Neurodegenerative Diseases of the Ministry of Education, Beijing 100069, China; Department of Functional Neurosurgery, Beijing Tiantan Hospital, Capital Medical University, Beijing 100070, China; Department of Functional Neurosurgery, Beijing Neurosurgical Institute, Capital Medical University, Beijing 100070, China; State Key Laboratory of Stem Cell and Reproductive Biology, Institute of Zoology, Chinese Academy of Sciences, Beijing 100101, China; Institute for Stem Cell and Regeneration, Chinese Academy of Sciences, Beijing 100101, China; National Stem Cell Resource Center, Chinese Academy of Sciences, Beijing 100101, China; University of Chinese Academy of Sciences, Beijing 100049, China

**Keywords:** Ephrins, NCK2, Parkinson’s disease, SNCA, Synucleinopathies

## Abstract

Synucleinopathies such as Parkinson’s disease, dementia with Lewy bodies and multiple system atrophy are characteristic for **α**-synuclein aggregates in neurons or glia, and are always manifested olfaction deficits at their primary onsets. It remains elusive why aggregation of **α**-synuclein predominantly affect the olfactory system. Employing the knockout mice, we investigate the physiological function of α-synuclein in olfactory system. We found that deletion of α-synuclein primarily interferes the projection of olfactory sensory neurons. iTRAQ based LC-MS identified that 188 proteins are differentially expressed, including 9 that were associated with axon guidance. Among them, NCK2 is most significantly down-regulated, which was indicated to be involved a PPI network of 21 proteins, including 11 players of the Ephrin receptor signaling pathway. Either α-synuclein deletion or NCK2 deficiency can inactivate Eph A4 receptor. Re-expressing α-synuclein in the α-synuclein knockout neurons reverse the NCK2, as well as the phosphorylated Eph A4 (the activated Eph A4). Thus, α-synuclein regulates axon guidance through NCK2-Eph A4 signaling pathway. Malfunction of α-synuclein, whether because of deletion or aggregation, may cause aberrant olfactory neurons projection and subsequent olfaction deficits. This extended our knowledge of effects of α-synuclein in olfactory system, which may explain why olfaction is usually impaired in some synucleinopathy related disorders such as Parkinson’s disease.

---

Parkinson’s disease (PD) is the second most common neurodegenerative disorder with a prevalence of 0.5%-1% among persons 65–69 years of age and of 1%-3% among those over the age of 80 years **[1]**. Its pathogenesis remains elusive. However, it was widely accepted that both environmental and genetic factors play roles in this disease**[2, 3]**. Studies of genes responsible for familial Parkinsonism/PD may yield critical insights into mechanisms shared by sporadic and familial disease.

The first identified genetic mutation linked to heritable PD were found in SNCA, also known as α-synuclein (α-syn) which is responsible for an autosomal-dominant form of PD. α-syn is a small, well-conserved protein that is expressed in many tissues and cell types **[4–6]**; its main protein domains comprise an amphipathic region, non-amyloid-b component (NAC) domain and an acidic tail**[7–9]**. Several dominant-inherited single point mutations in SNCA have been identified in families that develop early onset PD**[10–15]**. Nonetheless, multiplications of the gene locus **[16,17]** and polymorphisms are associated with high risk of developing PD **[18,19].** Of note, α-syn not only contributes to familial Parkinsonism, it also has been shown to have central pathogenic role in sporadic PD **[20,21]**. Accumulated α-syn is the main component of Lewy bodies (LBs), the hallmark of PD. Hence, α-syn remains the most potent culprit underlying PD. Yet both pathogenic mechanisms underlying α-syn with PD and its physiological function are not clear.

The characteristic symptoms of PD include motor deficits such as rigidity, bradykinesia, postural instability, and a tremor at rest **[22–23]**, resulting from the progressive loss of dopaminergic neurons in the pars compacta of the substantia nigra (SN) of the midbrain and consequent decrease in striatal dopamine levels **[24]**. However, The onset of PD is considered to commence at least 20 years prior to detectable motor abnormalities. This period is referred to as the prodromal phase where patients experience a range of non-motor symptoms such as olfactory dysfunction, sleep disturbances, obesity and depression. Among all above symptoms, olfactory dysfunction is one of the first manifestations of PD. It is reported to precede motor symptoms by 20 years **[25,26]**. It has been shown a high prevalence of olfactory dysfunction in PD patients (45% to 90%). Braak and colleagues found□ □α-syn accumulated in olfactory bulb in the very early stage of PD. In his theory, an external pathogen accesses olfactory and transit to the olfactory bulb which might induce pathology in the above initiation site **[27,28]**. Then study the physiological function of α-syn in olfactory system will be helpful for us to understand the occurrence and development of PD.

Here, we use SNCA knockout mice and its littermate control to study the influence of α-syn deficiency on olfactory system. The α-syn knockout mice exhibited aberrant projection of olfactory sensory neurons (OSNs) and olfaction impairment accordingly. NCK2 identified by iTRAQ based LC-MS technique, and its downstream EphA4 were shown to be responsible for this α-syn efficiency-induced aberrant projection.

## Results

### α-syn was highly expressed in both olfactory epithelium and olfactory bulb

In order to investigate **α**-syn expression pattern in the brain, protein from different brain region including cortex (Crt), cerebellum (Cere), olfactory bulb (OB), olfactory epithelium (OE), striatum (Str) and substantia nigra (SN) was extracted and western blot was used to evaluate **α**-syn protein level. As shown in Fig 1, **α**-syn was detected in various brain regions. It was abundant in olfactory system, especially in the olfactory bulb. Immunofluorescence staining showed that α-syn is specifically expressed in the olfactory receptor neurons and the olfactory bundles of OE and the expression of α-syn was mainly focused on olfactory glomeruli and outer plexiform layer of OB. (Fig 1)

**Fig. 1.**
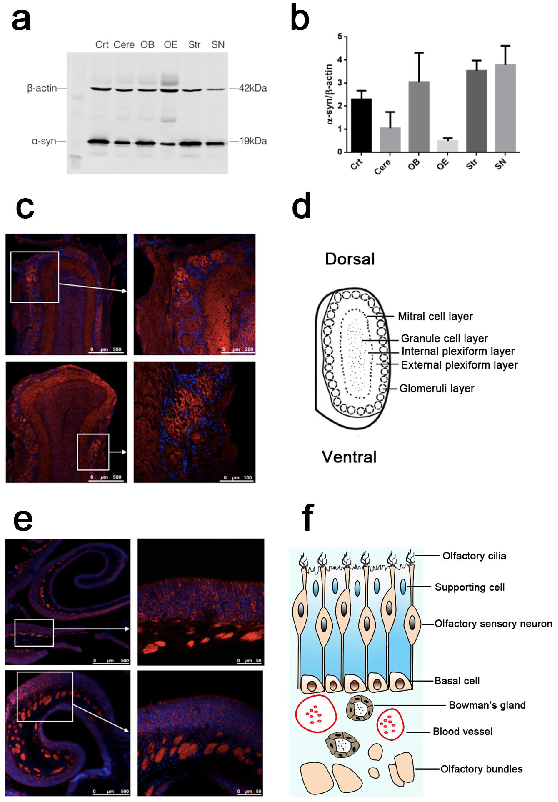
α-syn was highly expressed in both olfactory epithelium and olfactory bulb. (a-b) Western blotting to show α-syn expression pattern in different brain regions. α-syn was detected in various brain regions. It was abundant in olfactory system, especially in the olfactory bulb. Data are expressed as mean±SD, N=3 (c) Immunofluorescence staining to show α-syn expression pattern in OB. The expression of α-syn was mainly focused on olfactory glomeruli and outer plexiform layer of OB. (d) Schematic diagram of OB. (e) Immunofluorescence staining to show α-syn expression pattern in OE. α-syn is specifically expressed in the OSNs and the olfactory bundles of OE. (f) Schematic diagram of OE.

### α-syn deficiency induced aberrent projection of olfactory neurons

As we know, the mapping of olfactory sensory neurons (OSNs) onto glomeruli astonishingly precise. Each glomerulus receives axons from a large number of OSNs that express the same olfactory receptor (OR). So doing immunostainning of OB with one specific olfactory receptor antibody, only one positive glomerulus in each bulb will be detected. Olfr1507, one kind of OR, antibody was used to evaluate the axon projection in α-syn knockout mice in the present study. Of note, Olfr1507 positive axons projected aberrantly in α-syn deficient mice. Some Olfr1507 positive axons projected to external plexiform layer. Some projected to the inner plexiform layer. And some projected out of the glomerulus although it still locates in the glomerulus layer. (Fig 2) Buried pellet test and two-bottle preference test were used to evaluate the olfactory function of α-syn KO mice. In the buried food test, KO mice spent more time to locate the buried pellet compared with WT mice by 6 months. (P<0.01, Fig 2). In the two-bottle preference test, KO mice showed an increased intake of water from the bottle which tube was treated with 0.7% hydrochloride acid, in comparison to the control group by 6 months and 9 months. (P<0.05 for 6 months and P<0.01 for 9 months, two way ANOVA) In the two-bottle preference test, KO mice showed an increased intake of water from the bottle which tube was treated with 0.7% hydrochloride acid, in comparison to the control group. There were no significant differences between the strains in water intake before 3 months age, control animals still displayed a strong preference for water from non-treated bottle, however, the littermate KO mice started to consume water from bottle treated with 0.7% hydrochloride acid at 6 months old and kept increasing at 9 months old (P<0.05 for 6 months and P<0.01 for 9 months, Fig 2).

**Fig. 2.**
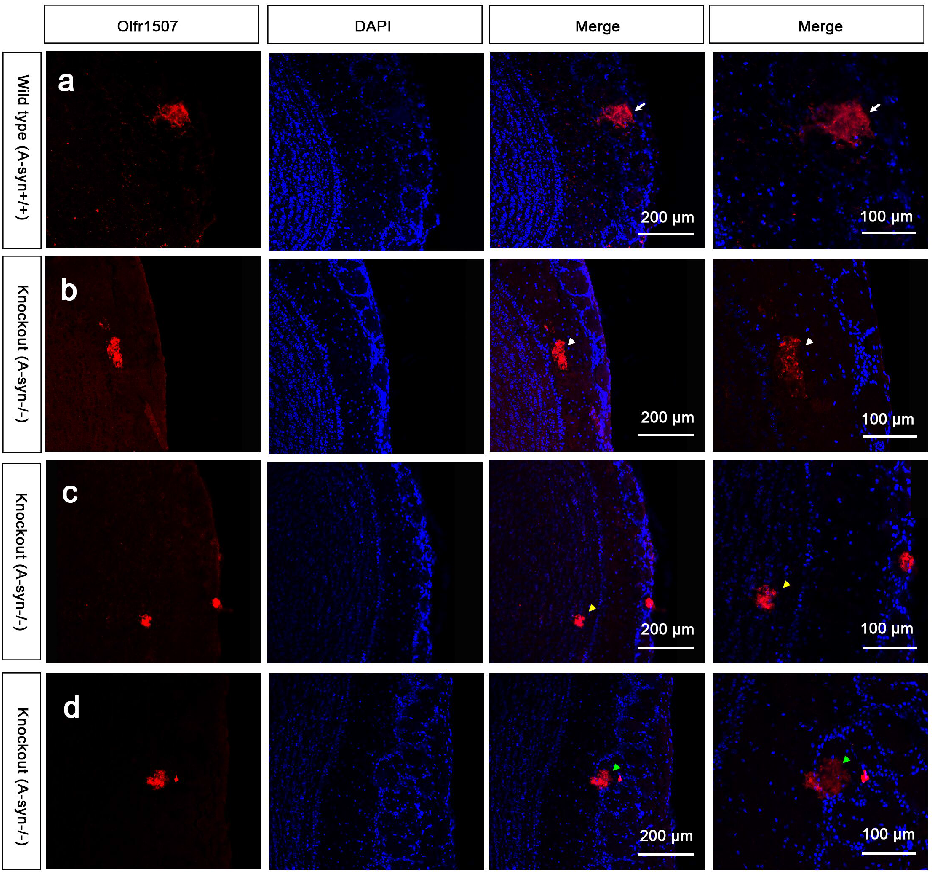
α-syn deficiency induced aberrant projection of olfactory neurons. (a) Immunostaining using an antibody against the Olfr1507 revealed that Olfr1507-expressing OSNs axon fibers converge to one specific glomerulus in each half bulb in wild type mice(WT A-Syn^+/+^) as indicated by the white arrow. In α-syn null mice (KO A-Syn^−/−^), Olfr1507-expressing OSNs axons projected aberrantly. Some Olfr1507 positive axons projected to external plexiform layer as indicated by the white arrowhead. Some projected to the inner plexiform layer as indicated by the yellow arrowhead. And some projected out of the glomerulus although it still locates in the glomerulus layer as indicated by the green arrowhead. (b) Statistical results of buried food pellet test. Data are expressed as the mean ±SD.**p < 0.01, ***p < 0.001, n=8. (c) Statistical results of two-bottle preference test. Data are expressed as the mean ±SD. *p < 0.05, **p < 0.01, n=6.

### 9 axon guidance associated proteins was identified by iTraq based LC-MS

In α-syn knockout mice, Olfr1507 positive axons project aberrantly which indicates that α-syn played an important role in olfactory axons guidance. To find out the molecules involved in this process, iTraq based LC-MS was carried out to compare the differentially expressed proteins between α-syn knockout mice and its litter mate control.

The screening criteria of reliable proteins: Unique peptide ≥ 1, remove invalid values and the anti-library data, and screening significant differentially expressed protein based on the reliable proteins. The screening criteria of differential protein: Fold change is 1.5. Multiple changes greater than 1.5 or less than 0.67 are considered to be differential proteins. Screening results statistics of differential proteins were as follows:

1. Total number of all proteins: 5489 pcs.

2. Total number of reliable proteins: 4936 pcs.

3. Number of differential proteins in Dynamics: 278 pcs.

4. The number of differential proteins in α-syn knockout mice and its litter mate control: 188 pcs.

Among all above 188 differential proteins, 133 were up-regulated and 55 were down-regulated. (Fig 3) Further analysis showed that 9 of differential proteins were associated with axon guidance. (Table I) The expression level of these 9 differential proteins was first verified by quantative realtime RT-PCR. The results showed that 5 of them including Cdh4, Cldn5, Dscam, Eif2b2, Pllp, Snapin were up-regulated and 4 of them including Snapin, Spag1, Stk11 and NCK2 were down-regulated. NCK2 was one of the most significant one that down-regulated by α-syn deletion. (data not shown)

**Table I.**
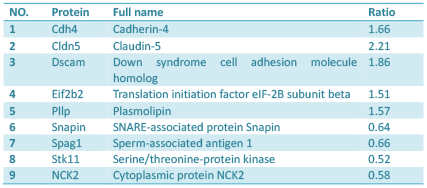
Axon guidance associated differential proteins.

**Fig. 3.**
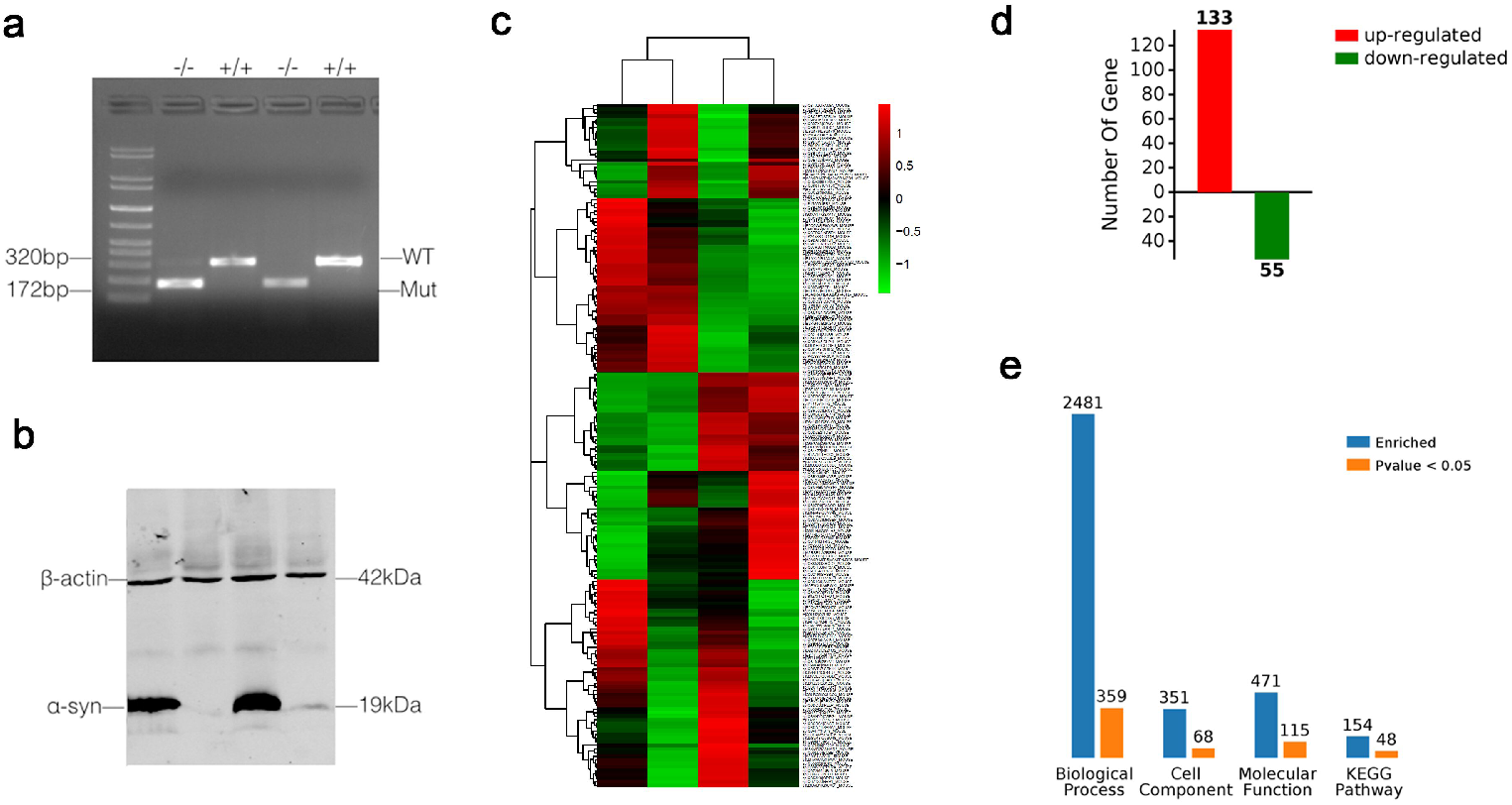
Identification of differentially expressed proteins in α-syn null mice by iTRAQ based LC-MS. (a-b) Genotyping and western blotting to to confirm the mice genotype; (c) The heatmap for the differentially expressed proteins. In the heatmap, red color represents up-regulated differentially expressed proteins and the green color represents down-regulated differentially expressed proteins. (d) statistical summary for differentially expressed proteins. A total of 188 proteins, can be seen in the figure in the difference comparison group. There are 133 pcs of up-regulation (red) of the protein and 55 pcs of down-regulation (green) of the protein. (e) All the differences in protein GO enrichment (BP, CC, MF) and pathway KEGG enrichment results and significant (p < 0.05) number were summarized. The blue column represents the total number of enrichment, orange column represents a significant number of enrichment. A protein is usually involved in a number of functions or pathways, so the total number of GO enrichment and pathway KEGG enrichment results will be far greater than the number of differential proteins. The significant function or the pathway means that the difference of the enrichment to the function or the pathway is significant.

### α-syn regulated NCK2 protein expression level

Cytoplasmic protein NCK2, one of the most significant down-regulated proteins, is shown to bind and recruit various proteins involved in the regulation of receptor protein tyrosine kinases family(RTKs). NCK2 is believed to be involved in axon guidance through the regulatory activities on Eph/Ephrin family, one of the important RTKs. As shown above, NCK2 is also one of the most significantly down-regulated proteins in **α**-syn knockout mice in the present study. (Table I) This down-regulation was further confirmed both in mRNA level and protein level by realtime RT-PCR and western-blotting. Nonetheless, the NCK2 down-regulation was reversed by overexpression of **α**-syn in the **α**-syn^−/−^ primary cultured neurons. (Fig 4) On the other side, **α**-syn overexpression also induced NCK2 up-regulation in protein level. (Fig 4)

**Fig. 4.**
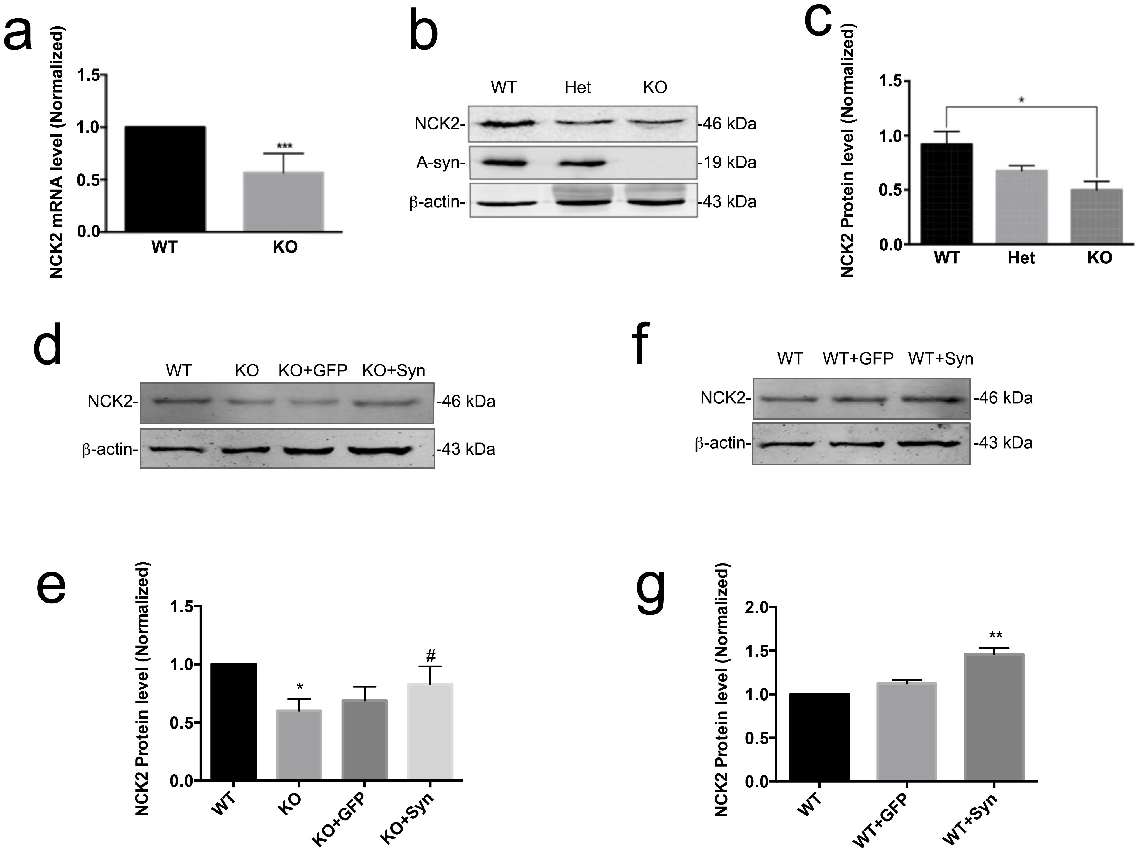
α-syn regulated NCK2 protein expression level. (a) NCK2 down-regulation was further confirmed in mRNA level by realtime RT-PCR. (b-c) NCK2 down-regulation was further confirmed in protein level by western-blot. (d-e) α-syn rescue experiment. NCK2 down-regulation was reversed by overexpression of **α**-syn in the α-syn^−/−^ primary cultured neurons. (f-g) α-syn overexpression also induced NCK2 up-regulation. Data are expressed as mean±SD. *p<0.05, **p<0.01 compared to WT; #p<0.05 compared to KO; n=4

### NCK2-EphA4 was involved in the mis-guidance of olfactory sensory neurons

Bioinformatics analysis showed that 21 proteins were enriched in PPI network with NCK2. (p<1.0e-16) (Fig 5) Biological analysis for these 21 proteins showed that 11 of them were enriched in Ephrin receptor signaling pathway. Molecular function analysis showed that 8 out of 21 proteins exhibited Ephrin receptor activity. KEGG analysis suggested that 16 were enriched in axon guidance pathway. (Table II) The axon guidance pathway showed that both Ephrin A and Ephrin B receptor signaling pathway involved in axon guidance, either on repulsion or attraction by cooperation with NCK2. (Fig 6) Further analysis was carried out on the association of NCK2 and all Ephrin receptors. The direct interactions between NCK2 and EphrinB1 or EphrinB2 or EphrinB3 or EphrinB4 or EphrinA1 or Ephrin A4 or EphrinA5 was experimentally confirmed. And indirect or predicted interactions also existed in NCK2 and Ephrin A2 or EphrinA3. (Fig 6)

**Table II.**
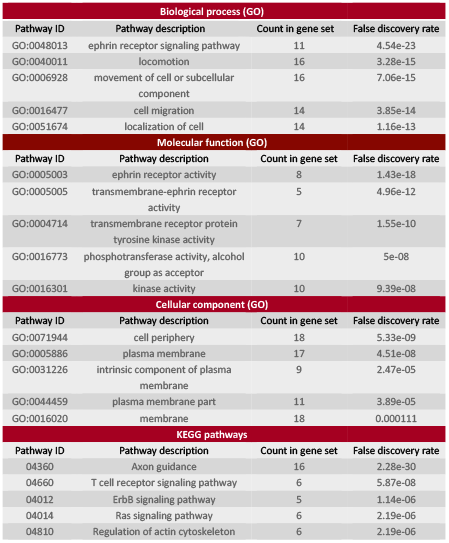
Bioinformatics analysis on 21 NCK2 associated proteins.

| Biological process (GO) |  |  |  |
| --- | --- | --- | --- |
| Pathway ID | Pathway description | Count in gene set | False discovery rate |
| GO:0048013 | ephrin receptor signaling pathway | 11 | 4.54e-23 |
| GO:0040011 | locomotion | 16 | 3.28e-15 |
| GO:0006928 | movement of cell or subcellular component | 16 | 7.06e-15 |
| GO:0016477 | cell migration | 14 | 3.85e-14 |
| GO:0051674 | localization of cell | 14 | 1.16e-13 |
| Molecular function (GO) |  |  |  |
| Pathway ID | Pathway description | Count in gene set | False discovery rate |
| GO:0005003 | ephrin receptor activity | 8 | 1.43e-18 |
| GO:0005005 | transmembrane-ephrin receptor activity | 5 | 4.96e-12 |
| GO:0004714 | transmembrane receptor protein tyrosine kinase activity | 7 | 1.55e-10 |
| GO:0016773 | phosphotransferase activity, alcohol group as acceptor | 10 | 5e-08 |
| GO:0016301 | kinase activity | 10 | 9.39e-08 |
| Cellular component (GO) |  |  |  |
| Pathway ID | Pathway description | Count in gene set | False discovery rate |
| GO:0071944 | cell periphery | 18 | 5.33e-09 |
| GO:0005886 | plasma membrane | 17 | 4.51e-08 |
| GO:0031226 | intrinsic component of plasma membrane | 9 | 2.47e-05 |
| GO:0044459 | plasma membrane part | 11 | 3.89e-05 |
| GO:0016020 | membrane | 18 | 0.000111 |
| KEGG pathways |  |  |  |
| Pathway ID | Pathway description | Count in gene set | False discovery rate |
| 04360 | Axon guidance | 16 | 2.28e-30 |
| 04660 | T cell receptor signaling pathway | 6 | 5.87e-08 |
| 04012 | ErbB signaling pathway | 5 | 1.14e-06 |
| 04014 | Ras signaling pathway | 6 | 2.19e-06 |
| 04810 | Regulation of actin cytoskeleton | 6 | 2.19e-06 |

**Fig. 5.**
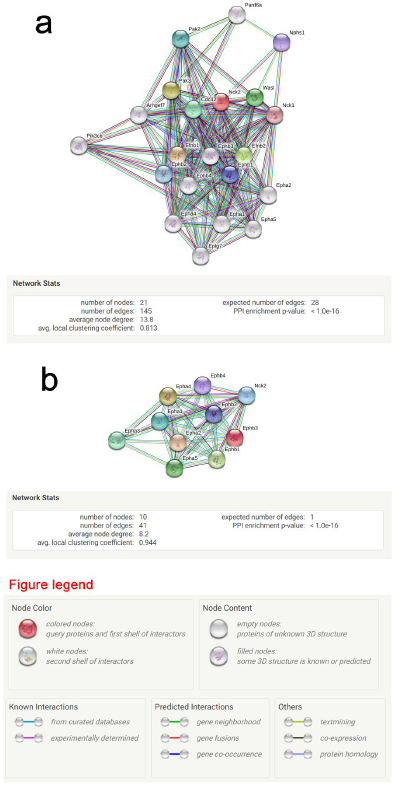
Bioinformatics analysis showed that 21 proteins were enriched in PPI network with NCK2. (a) PPI network with NCK2. Bioinformatics analysis showed that 21 proteins were enriched in PPI network with NCK2. Biological analysis for these 21 proteins showed that 11 of them were enriched in Ephrin receptor signaling pathway. (b) Further analysis on the association of NCK2 and all Ephrin receptors. And indirect or predicted interactions existed in NCK2 and Eph A1-A5, EphB1-B4.

**Fig. 6.**
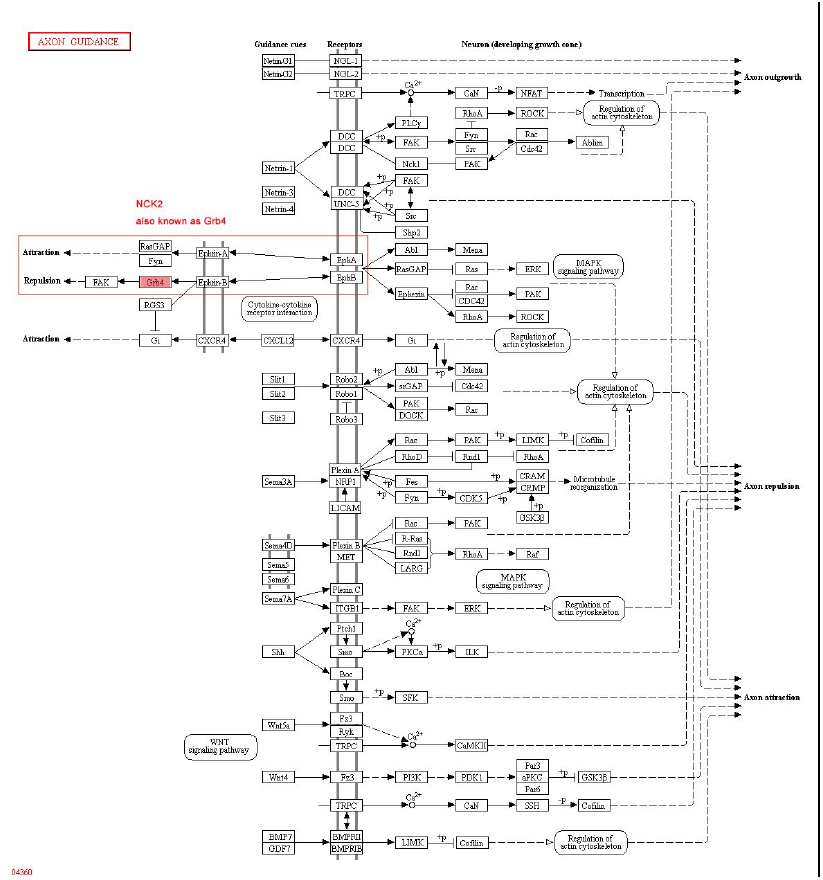
Axon guidance pathway analysis. The axon guidance pathway showed that both Ephrin A and Ephrin B receptor signaling pathway involved in axon guidance, either on repulsion or attraction by cooperation with NCK2.

To find out whether Ephrin receptor signaling pathway involved in the aberrant axon projection of olfactory sensory neurons induced by **α**-syn deficiency, all 9 phosphorylated Ephrin receptors were detected by western blot. As shown in Fig 7, EphA4 was inactivated which was indicated by its lower phosphorylated level in **α**-syn deficiency mice. Moreover, knocking-down NCK2 led to EphA4 inactivation as well. (Fig 7)

**Fig. 7.**
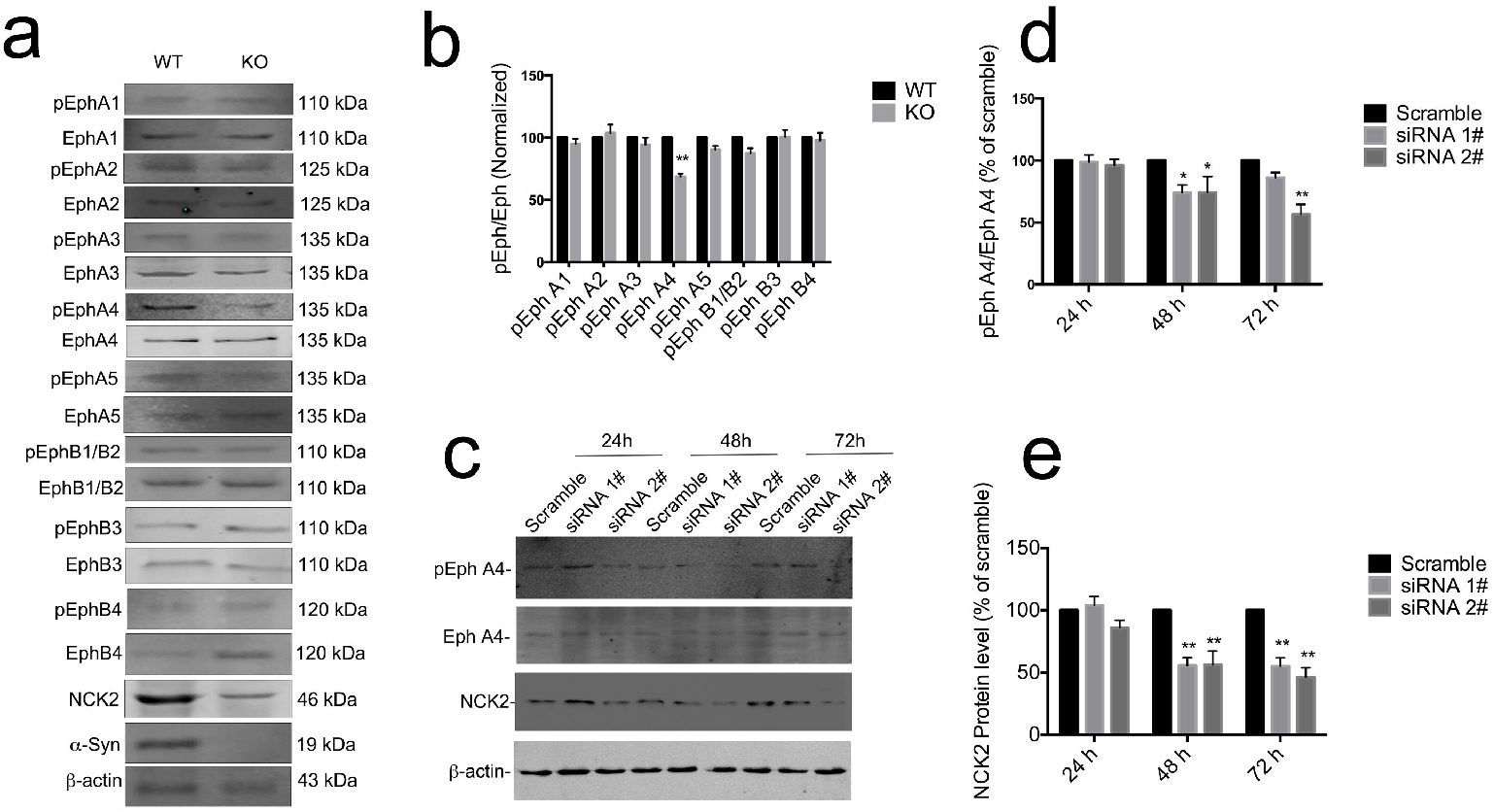
EphA4 was inactivated with α-syn or NCK2 deficiency. all 9 phosphorylated Ephrin receptors were detected by western blot. As shown in, Eph A4 was inactivated indicated by lower phosphorylated level in **α**-syn deficiency mice. However, the phosphorylation level of EphA1, EphA2, EphA3, EphA5, EphB1/B2, EphB3 and EhpB4 were not changed. (a-b) However, phosphorylated Ephrin A4 level was downregulated with NCK2 deficiency.(c-d) Data are expressed as mean±SD. *p<0.05, **p<0.01 compared to WT or scramble, respectively; n=3

**Fig. 8.**
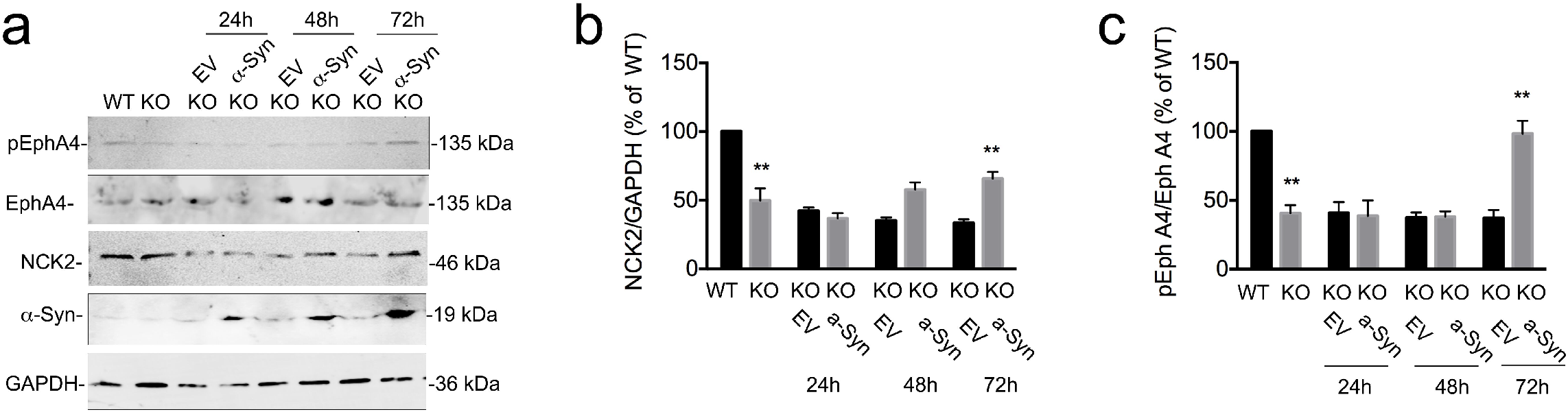
NCK2-EphA4 deactivation resulted from □-syn deficiency can be reversed by overexpression of □-syn. The primary cultured □-syn^/^ neurons were transfected with LV-□-syn. NCK2-EphA4 pathway was evaluated by western blot. As shown above, NCK2 and phosphorylated EphA4 were dow · n-regulated in □-syn^−/−^ neurons, indicating that NCK2-EphA4 pathway was deactivated. However, the downregulation of NCK2 and phosphorylated EphA4 were reversed by overexpression of □-syn.

To further confirm that NCK2-EphA4 was modulated by **α**-syn, the primary cultured **α**-syn^−/−^ neurons were transfected with LV-**α**-syn. NCK2-EphA4 pathway was evaluated by western blot. As shown above, NCK2 and phosphorylated EphA4 were down-regulated in **α**-syn^−/−^ neurons, (Fig 7) indicating that NCK2-EphA4 pathway was deactivated. However, the downregulation of NCK2 and phosphorylated EphA4 were reversed by overexpression of α-syn.

## Discussion

It was reported that **α**-syn null mice display striking resistance to MPTP-induced degeneration of DA neurons and DA release, and this resistance appears to result from an inability of the toxin to inhibit complex I **[29]**. But later Abeliovich et al showed that mice lacking **α**-syn displayed a reduction in striatal DA and an attenuation of DA-dependent locomotor response to amphetamine**[30]**. These findings support the hypothesis that α-Syn is an essential protein in central nervous system. It plays an important role in many physiological functions in vivo. Up to now, the phsyiological function of α-syn that was reported include: 1) α-syn functions in synaptic neurotransmitter release by binding to synaptic vesicles and regulating synaptic vesicle pool organization. **[31,32]**. 2) α-syn, a presynaptic protein that plays a crucial role in dopamine compartmentalization in the striatum, is also involved in the compartmentalization of norepinephrine in the dentate gyrus**[33]**. 3) α-syn modulates brain glucose metabolism because α-syn deficiency leads to increased glyoxalase I expression and glycation stress**[34]**. 4) What’s more, it was shown that α-syn knockout mice had decreased striatal dopamine and altered dopamine release, uptake in response to electrical stimuli, which suggests that α-syn may have a role in dopamine neurotransmission**[35]**. And similarly Connor-Robson et al. reported that stabilization of the striatal DA level depends on the presence of α-syn and cannot be compensated by other family members. **[36]** α-syn modulates dopaminergic neuron development, DA reuptake and stabilizes striatal DA level **[35–37]**. 5) α-syn modulates microglial phenotype and neuroinflammation. **[38,39]**6) in Kokhan et al’s experiment, α-syn knockout mice have cognitive impairments**[40]**. α-syn knockout mice showed impaired spatial learning and working memory.7)Using co-immunoprecipitation techniques, Alim et al found α-syn significantly interacts with tubulin and that α-syn may behave as a potential microtubule-associated protein **[41]**. Here we report a novel function of a-syn that it is essential for OSNs axon projection. α-syn was highly expressed in both olfactory epithelium and olfactory bulb. Lacking of α-syn induced aberrant projection of OSNs.

188 differentially expressed proteins resulted from α-syn deficiency were identified by iTraq based LC-MS. 9 of them were associated with axon guidance, including Cdh4, Cldn5, Dscam, Eif2b2, Pllp, Snapin, Spag1, Stk11, and NCK2. These results suggest that Cdh4, Cldn5, Dscam, Eif2b2, Pllp, Snapin, Spag1, Stk11, NCK2 may be involved in the modulation of olfactory neurons projection by α-syn. In the present study, we further confirmed that NCK2 was down-regulated in α-syn^−/−^ mice both in mRNA and protein level. NCK2 belongs to a family of adaptor proteins and NCK2 is located in chromosome 2. The protein contains three SH3 domains and one SH2 domain. The protein has no known catalytic function but has been shown to bind and recruit various proteins involved in the regulation of receptor protein tyrosine kinases. It is through these regulatory activities that this protein is believed to be involved in cytoskeletal reorganization. In the present study, we examined NCK2 targeted receptor protein tyrosine kinases Ephrin family and found that EphA4 was deactivated in α- syn^−/−^ mice. Down-regulation of NCK2 and deactivation of Eph4 can be reversed by overexpression of α-syn, indicating that NCK2-Eph A4 signaling pathway was responsible, at least partially responsible for this α-syn mediated aberrant projection.

In conclusions, α-syn regulates the axon guidance of olfactory neurons via NCK2-EphA4 pathway. The present findings provide an insight into α-syn physiological function. And NCK2 and EphA4 are the most important molecules that involved in the molecular mechanism of α-syn mediated axon guidance.

## Methods

### Animals

We used 8-week-old male α-syn KO mice and their male wild-type (WT) littermates for the present i-Traq based LC-MS study. α-syn KO and WT mice were group housed in a room with a 12 h light/dark cycle (lights on at 7:00 a.m.), with access to food and water ad libitum. Room temperature was kept at 23 ± 2 °C. All procedures were approved by the Institutional Animal Care and Use Committee of Capital Medical University.

### Western blot

Equal amounts of protein (100 μg) were separated by 10% sodium dodecyl sulfate polyacrylamide gel electrophoresis and then transferred to a PVDF membrane (Pall Co., Taiwan) that was blocked in 5% non-fat milk for 1 h at room temperature and incubated overnight with primary antibodies against the following proteins: α-synuclein (1:1000, BD), NCK2 (1:1000, Sigma-Aldrich), pEphA1 (1:500, Abcam), pEphA2(1:500, Abcam), pEphA3(1:500, Abcam), pEphA4 (1:500, Abcam), pEphA5 (1:500, Abcam), pEph B1/B2 (1:500, Abcam) and β-actin (1:1000, Sigma-Aldrich). Immunoreactivity was detected with Fluorescence-conjugated anti-mouse or -rabbit antibodies (1:10,000; Sigma-Aldrich).

### Immunohistochemistory

Mice were sacrificed and brains were fixed with 4% paraformaldehyde and sectioned at a 50-μm thickness. The staining was carried out on free-floating sections. Sections were rinsed in PBS and incubated in 3% H_2_O_2_ for 20 min to block the endogenous peroxidase activity. After washing in PBS, sections were incubated in blocking serum (10% goat serum and 0.1% Triton X-100 in PBS) for 30 min, followed by incubation in anti-□ □-synuclein antibody solution (1:1000, BD) or olfr1507 (1: 1000, Thermo Fisher) for 24 h at 4 ¤. Then, the sections were incubated with Alexa 594-conjugated goat anti-mouse IgG (1: 1000, Cell Signaling Technology) for 2h at room temperature. Images were captured by a Leica confocal microscope (Leica Microsystems, USA).

### Quantitative RT PCR

Total RNA was isolated from the olfactory bulb of α-syn KO or its littermate control using the RNeasy Mini kit (Qiagen). 5 μg RNA were reverse-transcribed in a reaction containing 1 μl dNTP (10 mmol/l; Invitrogen), 0.5 μg oligo(dT) primer, with 50 U Superscript RNase reverse transcriptase (Invitrogen) for 50 min at 42°C. Forward and reverse primer sequences for PCR were as follows: β-actin, 5’-ACC ACC ATG TAC CCA GGC ATT-3’ and 5’-CCA CAC AGA GTA CTT GCG CTC A-3’; *NCK2*, 5’-GTCATAGCCAAGTGGGACTACA-3’ *and* 5’-GCACGTAGCCTGTCCTGTT-3’. The reaction was carried out on 7500 real-time PCR systems (Applied Biosystems, Foster City, CA, USA) using the default thermal cycling conditions (2 min at 50°C plus 10 min at 95°C for the hot start; and 40 cycles of 15 s at 95°C plus 1 min at 60°C).

### iTRAQ based LC-MS

OB tissue were taken from 8-week-old male α-syn KO mice and their male wild-type littermates. Two biological replicates of each group were prepared for the proteomics experiments. Briefly, the total protein of OB was grinded and dissolved in lysis buffer [9 M Urea, 4% CHAPS, 1%DTT, 1%IPG buffer (GE Healthcare)]. The mix was incubated at 30°C for 1 hour and centrifuged at 15,000 g for 15 min at room temperature. The supernatant was collected and quantified by the Bradford method.

100 μg of protein of each sample was dissolved in a dissolution buffer (AB Sciex, Foster City, CA, USA). After being reduced, alkylated and trypsin-digested, the samples were labeled following the manufacturer’s instructions for the iTRAQ Reagents 8-plex kit (AB Sciex). Samples were each labeled with iTRAQ reagents with molecular masses of 113, 114, 115, and 116 Da, respectively. After labeling, all samples were pooled and purified using a strong cation exchange chromatography (SCX) column by Agilent 1200 HPLC (Agilent) and separated by liquid chromatography (LC) using a Eksigent nanoLC-Ultra 2D system (AB SCIEX). The LC fractions were analyzed using a Triple TOF 5600 mass spectrometer (AB SCIEX). Mass spectrometer data acquisition was performed with a Triple TOF 5600 System (AB SCIEX, USA) fitted with a Nanospray III source (AB SCIEX, USA) and a pulled quartz tip as the emitter (New Objectives, USA). Data were acquired using an ion spray voltage of 2.5 kV, curtain gas of 30 PSI, nebulizer gas of 5 PSI, and an interface heater temperature of 150°C. For information dependent acquisition (IDA), survey scans were acquired in 250 ms and as many as 35 product ion scans were collected if they exceeded a threshold of 150 counts per second (counts/s) with a 2^+^ to 5^+^ charge-state. The total cycle time was fixed to 2.5 s. A rolling collision energy setting was applied to all precursor ions for collision-induced dissociation (CID). Dynamic exclusion was set for ½ of peak width (18 s), and the precursor was then refreshed off the exclusion list.

The iTRAQ data were processed with Protein Pilot Software v4.0 against the *A. japonicus* database using the Paragon algorithm. Protein identification was performed with the search option of emphasis on biological modifications. The database search parameters were the following: the instrument was TripleTOF 5600, iTRAQ quantification, cysteine modified with iodoacetamide; biological modifications were selected as ID focus, trypsin digestion. For false discovery rate (FDR) calculation, an automatic decoy database search strategy was employed to estimate FDR using the PSPEP (Proteomics System Performance Evaluation Pipeline Software, integrated in the ProteinPilot Software). The FDR was calculated as the number of false positive matches divided by the number of total matches. Then, the iTRAQ was chosen for protein quantification with unique peptides during the search, and peptides with global FDR values from fit less than 1% were considered for further analysis. Within each iTRAQ run, differentially expressed proteins were determined based on the ratios of differently labeled proteins and p-values provided by Protein Pilot; the p-values were generated by Protein Pilot using the peptides used to quantitate the respective protein. Finally, for differential expression analysis, fold change was calculated as the average ratio of 113/ 114, 113/116, 115/114 and 115/116. Proteins with a fold change of >1.5 or <0.67 and p < 0.05 were considered to be significantly differentially expressed.

### Quantification and statistical analysis

Data are expressed as mean ± SD. The sample size (n) represents biological replicates. Student’s t test was used for comparison of two group averages. When there were more than two groups, one-way ANOVA Tukey-Kramer multiple comparisons Bonferroni post hoc test was performed. All the bioinformatics were analyzed or confirmed by bioinformaticians. iTRAQ based LC-MS analysis was performed in a double-blinded manner. A P value <0.05 was considered statistically significant.

## ACKNOWLEDGEMENTS

This work was supported by grants from the National Natural Science Foundation of China (81830033), Natural Science Foundation of Beijing (7192017), Scientific Research Common Program of Beijing Municipal Commission of Education (KM201710025001).

